# Assessing SARS-CoV-2 spatial phylogenetic structure: Evidence from RNA and protein sequences

**DOI:** 10.1101/2020.06.05.135954

**Authors:** Leandro R. Jones, Julieta M. Manrique

## Abstract

Severe acute respiratory syndrome coronavirus 2 (SARS-CoV-2) is an emergent RNA virus that spread around the planet in about 4 months. The consequences of this rapid dispersion are under investigation. In this work, we analyzed thousands of genomes and protein sequences from Africa, America, Asia, Europe, and Oceania. We show that the virus is a complex of slightly different variants that are unevenly distributed on Earth, and demonstrate that SARS-CoV-2 phylogeny is spatially structured. Remarkably, the virus phylogeographic patterns were associated with ancestral amino acidic mutations. We hypothesize that geographic structuring is the result of founder effects occurring as a consequence of, and local evolution occurring after, long-distance dispersal. Based on previous studies, the possibility that this could significantly affect the virus phenotype is not remote.

## Introduction

A cluster of acute atypical pneumonia syndrome cases of unknown etiology was reported late December 2019 in China. Rapidly, metagenomic RNA sequencing of bronchoalveolar lavage fluid showed that the etiological agent was a new human coronavirus (Wu et al., 2020; Zhou et al., 2020), now named severe acute respiratory syndrome coronavirus 2 (SARS-CoV-2; Gorbalenya et al., 2020). Human coronaviruses are well known to produce mild to moderate upper-respiratory tract illnesses. However, severe acute respiratory syndrome coronavirus (SARS-CoV) responsible for a 2002-2003 epidemic, Middle East respiratory syndrome coronavirus (MERS-CoV) responsible for a 2012 outbreak and the novel SARS-CoV-2, cause severe syndromes with high fatality rates (10%, 36% and 4 %, respectively) (Cui et al., 2019; WHO, 2020). SARS-CoV and SARS-CoV-2 are ~79% identical at the genetic level and share important features as the same cell receptor and immunopathogenic mechanism.

SARS-CoV-2 has dispersed to hundreds of countries, causing millions of infections. The consequences of this rapid spread on the virus evolution are still unclear. Recent studies suggested that SARS-CoV-2 genotypes are heterogeneously distributed (Forster et al., 2020; Rambaut et al., 2020). However, quantitative and phenotypic assessments are still lacking. When relatedness between spatially coexisting lineages is greater than expected, their distribution is said to be phylogenetically structured (Webb et al., 2002). One of the consequences of structuring is spatially close sequences being more similar to each other than expected by chance (Figure S1). The phenomenon can be assessed by comparing measured phylogenetic distances with those expected under no structuring, which can be accomplished by Monte Carlo simulations. Here, we describe an analysis of SARS-CoV-2 genomes collected from the beginning of the pandemic to April 25, 2020. We observed an uneven distribution of genetic and amino acidic variants, and a statistically significant spatial phylogenetic structure. Notably, ancestral amino acidic substitutions were highly fitted to the virus phylogeny and geographic patterns, strongly suggesting that long-distance dispersal can facilitate the establishment of otherwise rare, and/or the emergence of new, viral phenotypes.

## Materials and Methods

### Operational Taxonomic Units delimitation and Datasets

Structure analyses require to determine the abundance patterns of a set of operational taxonomic units (Webb et al., 2002). We initially studied a small dataset of 831 genomes. The smallness of this sample allowed us to identify sequence variants (SV) from all the possible pair-wise Needleman-Wunsch alignments in the dataset. By the other side, we compiled a much larger dataset (n=8,612 genomes), aiming to gather a wider genomes sample to assess the virus spatial phylogenetic structure. The procedure used to delimit the small dataset SVs has a time complexity of *O(n*^*2*^*L)* − *O(n*^*3*^*L)*, where *n* and *L* are the sequences’ number and length, respectively. Thus, the approach could not be used with the large dataset, for which pair-wise distance were obtained from the sequences’ multiple alignment (details below). The small dataset studied here consisted in all the genomes described from the beginning of the pandemic to March 14, 2020 (Table S1). These sequences were first inspected for the presence of ambiguous and undetermined positions by the *base.freq()* function from the *Ape* package (Paradis and Schliep, 2018). The selected data (genomes with no non-DNA characters) were aligned with *Mafft* (Katoh and Toh, 2008) (details below) and the obtained alignment was visually inspected with *Jalview* (Waterhouse et al., 2009). This revealed the rare presence of sequences with internal and terminal gaps, which were dismissed (Table S1). After removing these sequences and unifying sequences recorded in both GenBank and GISAID’s EpiCoV™, we ended up with 353 genomes. From these, we generated 62,128 Needleman-Wunsch pairwise alignments and, for each one, counted the number of mismatches and indels. End gaps were excluded from the counts since these correspond to sequencing differences. Pairwise alignments and hamming distances were obtained with the *nwhamming()* function of the *Dada2* package (Callahan et al., 2016). Shortly, we made an R script that takes a sequence (the *query*) from a dataset, aligns it against the rest of sequences and calculates the corresponding hamming distances:

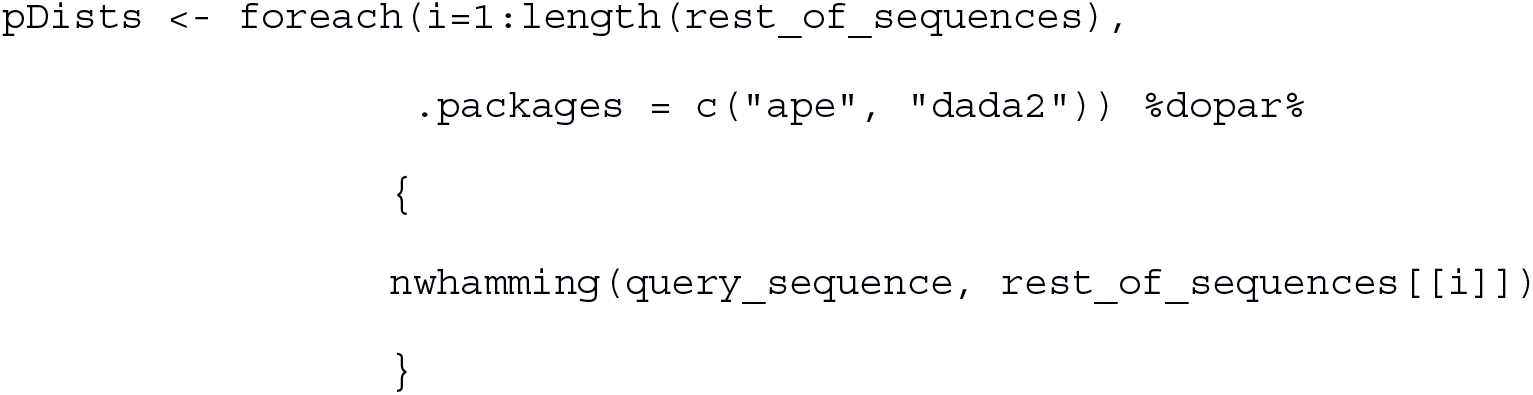

Based on the obtained distances, the script determines which sequences are identical to the query sequence, put these together in an identity cluster, and starts a new iteration. This was repeated until all the sequences in the dataset were processed. We ensured that the sequences representative of each SV used in posterior analyses were the longest of each identity cluster. The large dataset studied here consisted in all the sequences in GISAID’s EpiCoV™ Database sampled between December 2019 and April 25, 2020 presenting less than 1 % non-DNA characters (*high coverage only* option) (Elbe and Buckland-Merrett, 2017; Shu and McCauley, 2017). The corresponding sequences and metadata are described in Table S2. As a first step we filtered-out all the sequences presenting non-DNA characters. Previous to clustering and phylogenetic analyses, we aligned the sequences and evaluated the diversity along the obtained alignment by inspecting the entropy at each aligned position using the *HDMD* package (https://CRAN.R-project.org/package=HDMD). The alignment positions homologous to nucleotides 1-77 and 29,804-29,903 of the isolate Wuhan-Hu-1 (NC_045512.2) presented too high diversities compared to the rest of the genome (Fig. S2). Thus, these regions were masked-out in posterior analyses. Furthermore, the sequences presenting missing data after and before positions 77 and 28,804, respectively, and rare internal gaps were dismissed. The final, high-quality dataset had 4,333 genomes. Pairwise distances were obtained from the aligned sequences using R string comparison tools, and processed similarly to what was done with the small dataset.

### SV abundances

If the variants of a virus are randomly distributed among human populations, the probability of detecting a given SV *i*, in some area *A*_*j*_, *P(V*_*i*_|*A*_*j*_), should be the same for all *j*: *P(V*_*i*_|*A*_*1*_)=*P(V*_*i*_|*A*_*2*_)=…=*P(V*_*i*_|*A*_*N*_)=*P(V*_*i*_), where *N* is the number of sampled areas. We modeled the probability of observing a given SV, *P(V*_*i*_), by its frequency in the data. Then, the expected number of SV *i* sequences in region *A*_*j*_ can be obtained as *P(i)* × *M*_*j*_, where *M*_*j*_ is the number of sequences from *A*_*j*_. Standardized deviations from expectation can be obtained by dividing the squared deviations *(O*_*ij*_ − *E*_*ij*_)^*2*^ by *E*_*ij*_, where *O*_*ij*_ and *E*_*ij*_ are the observed and expected numbers of SV *i* sequences in region *j*, respectively. This statistic is not affected by the order of magnitude of the data, which allows to straightforwardly compare very abundant and rare SVs, and shallowly and much sampled regions.

### Phylogenetic analysis

Sequence alignments were obtained with *Mafft*’s *FFT-NS-2* algorithm. Under this strategy, the program first generates a rough alignment (namely *FFT-NS-1* alignment) from a guide tree created from a distance matrix made by computing the number of hexamers shared between all sequence pairs. Then the *FFT-NS-1* alignment is used to generate a new tree, which is used to guide a second progressive alignment. Once aligned, the small dataset presented 512 variable sites of which 109 were parsimony informative. The large dataset alignment presented 2,713 polymorphic sites of which 885 were parsimony informative. The small dataset was phylogenetically analyzed with *PhyML* version 3.3 (Guindon et al., 2010) under a generalized time-reversible model with a proportion of invariable sites and 4 rate categories. The nucleotide frequencies, proportion of invariable sites and value of the gamma shape parameter were maximum-likelihood estimated. Tree searches consisted in 100 parsimony starting trees that were refined by *NNI* and *SPR* branch swaps (option *-s BEST*). Branch supports were estimated with *PhyML*’s bootstrap routine. The large dataset analyses were performed with IQTREE (Minh et al., 2020) using the GTR+I+G model of nucleotide substitution (selected by the IQTREE’s BIC routine) with base frequencies inferred from the data and the default tree search algorithm. Branch supports were calculated by ultrafast bootstrap (Hoang et al., 2017) (n=1,000) in IQTREE. Trees were inspected and plotted by *Dendroscope* (Huson and Scornavacca, 2012), *Ape* and *ggtree* (Yu, 2020). Ancestral character states were inferred with the *ace()* function from Ape using the SVs’ sequences and phylogeny.

### Spatial phylogenetic structure

Structure analyses were performed by the *picante* package (Kembel et al., 2010). Null distributions were inferred by shuffling SVs across tree tips 10,000 times. Using the obtained permutations and the actual data, the software calculates the following weighted metric for each region:

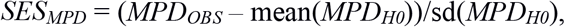

where *MPD*_*OBS*_ is the mean phylogenetic distance (*MPD*) between all sequences from the region, *mean(MPD*_*H0*_) is the average of the mean phylogenetic distance (*MPD*) between the sequences from the region in the randomized data and *sd(MPD*_*H0*_) is the corresponding standard deviation. Negative and low *SES*_*MPD*_ values support structured distributions. Significance levels are calculated as the *MPD*_*OBS*_ rank divided by the number of permutations minus one.

## Results

### SV analyses

Out of 62,128 pairwise comparisons performed from the small dataset, 61,759 (99.4%) revealed 1 or more mutations and 36,771 (59.1%) showed between 5 (Q1) and 11 (Q3) changes. The analysis of the large dataset also revealed a very large diversity, as depicted in Figure 1. Out of 9,385,278 genome pairs compared, only 50,839 (0.5%) were identical and 5,056,965 (53.8%) showed between 6 (Q1) and 12 (Q2) reciprocal mutations. The number of detected SVs relative to the total number of sequences analyzed was very similar for both datasets and methods. For the small dataset we observed one SV every 1.55 sequences, while for the large one the ratio was of 1.85. The small dataset presented 228 unique sequences or SVs. The most abundant SV was represented by 30 sequences. Twenty five were represented by 2 sequences each, 9 by 3 sequences, 3 by 4, 4 by 5 and the remaining three by 7, 9 and 14 sequences. These results are summarized in Table S3. The large dataset presented 2,336 operational taxonomic units or SVs. The most abundant SV was represented by 224 sequences. Eleven SVs presented more than 30 sequences each, 27 presented at least 10 sequences and 365 presented between 2 and 10 sequences. These results are detailed in Table S4.

**Figure 1.**
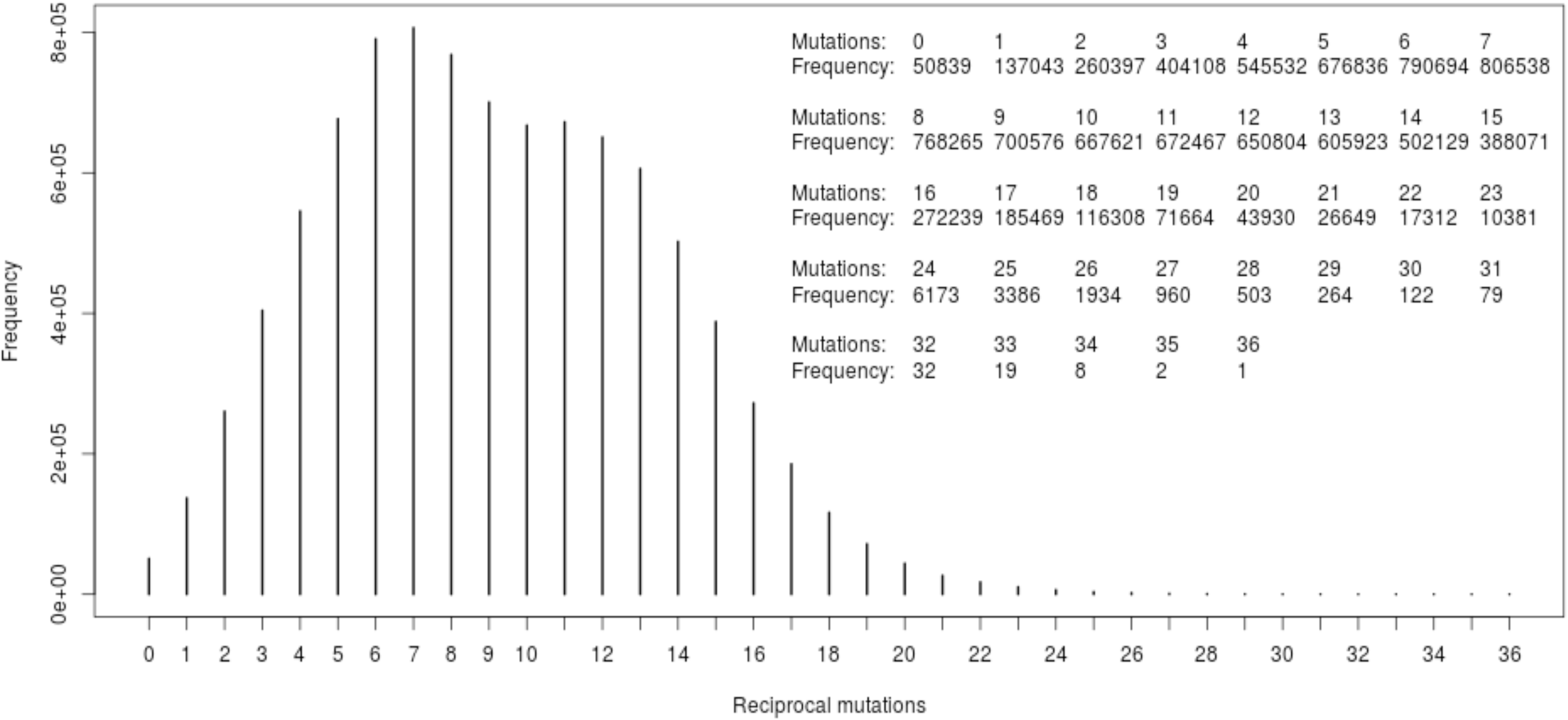
Pairwise genetic distances between the genomes from the large dataset studied here. The units of distance measures (reciprocal mutations) are nucleotide substitutions.

### SVs distribution

In the small dataset, the 24 most abundant SVs in Europe (n=78 genomes; SVs 2, 6, 7, 8, 10, 11, 16, 17, 18, 19, 20, 24, 29, 30, 34, 37, 38, 39, 40, 41, 42, 43, 44 and 45; Table S3) were absent from other continents and SV 1 was very prevalent in Asia (22 genomes) in comparison to North America (7 genomes). This can be easily appreciated from the deviations of the observed SV abundances from those expected if the virus variants were randomly distributed (Table S3). SVs 2, 6, 7, 8, 10, 11, 16, 17, 18, 19, 20, 24, 29, 30, 34, 37, 38, 39, 40, 41, 42, 43, 44 and 45 had a mean deviation from expectation of 2.4 in Europe, whereas, for the same region, the rest of SVs had a mean deviation of 0.68. Likewise, SV 1 deviations from expectation in Asia and Europe were 8.37 and 11.08, respectively, whereas in North America, Oceania and South America these were of only 2.26, 0.93 and 0.085, respectively. This reflects this SV's over-abundance in Asia and rarity in Europe, and that its frequencies in the rest of the continents are relatively well predicted by its global abundance. (Materials and Methods). The counts and distributions of the 2,336 SVs identified from the large dataset are detailed in Table S4. As observed for the small dataset, these SVs distribution was very uneven. Here we highlight the most relevant cases. Of the SVs represented by at least 10 sequences, only SVs 17 and 37 were distributed more or less homogeneously. The SV 1 abundance was smaller than expected in Asia, and larger than expected in North America. The SV 2 was represented only among the North American sequences and the SV 3 was over-abundant in Europe and scarcely represented in North America. The SVs 4, 9, 12, 23 and 29 were over-represented (SVs 4, 9 and 12) or exclusively observed (SVs 12 and 23) in Asia. The SVs 5 and 6 were too abundant in Europe and South America and were under-represented in Asia and North America. The SVs 7, 8, 10, 14, 16, 19, 20, 21, 24, 26, 30, 31, 33, 34, 35 and 38 were over-represented or exclusively detected in Europe. The SVs 11, 13, 15, 18, 22, 27 and 36 displayed more sequences than expected by chance in North America. The number of SV 28 genomes observed among the Oceania samples was about 20 times greater than expected. Likewise, the SVs 25 and 32 counts in South America and Africa, respectively, were approximately 22 and 21 times larger than the corresponding expected numbers.

### Phylogenetic structure analyses

The small dataset phylogeny is shown in Figure 2. A heat map describing the SVs distribution is shown alongside the phylogeny. Tree terminals are aligned with the heatmap cells, so that the Figure depicts the tree lineages’ abundances at each continent (heatmap columns). Clades *A*, *B*, H, *I* and *J* were distributed in Europe. Likewise, the majority of sequences located at clades *A*-*D*, and the sequences grouped in clades *F* and *G* were common in Europe. The exceptions were SVs 226, 46 and 112, which correspond to samples from North America (SVs 226 and 112) or from North America and Europe (SV 46). Most sequences in the branch containing clades *M* and *N* are from North America. Clades *E* and *K* sequences are all from Asia. As expected from the visual inspection of the SVs phylogeny and distribution, structuring was highly significant for North America and Europe (Table 1). The phylogeny and distributions of the large dataset SVs are detailed in Figure S3. A thorough inspection of these results revealed six major sections of the tree in which the sequences were clumped according to their origins, as shown in a simplified manner in Figure 3. For practical reasons, we refer to these six tree sections as *A*, *B*, *C*, *D*, *E*, and *F*. The North American sequences were more abundant in sections *A* and *E* while the European ones were prevalent in sections *B*, *C* and *F*, and moderately represented in section *A*. The Asian sequences clustered preferentially in section *D*, and in sections *C*, *E* and *F* but in proximal positions relative to the positions occupied by the North American (region *E*) and European (regions *C* and *F*) sequences. Limitation in space was significant (p < 0.01) for Asia, Europe and North America (Table 2).

**Figure 2.**
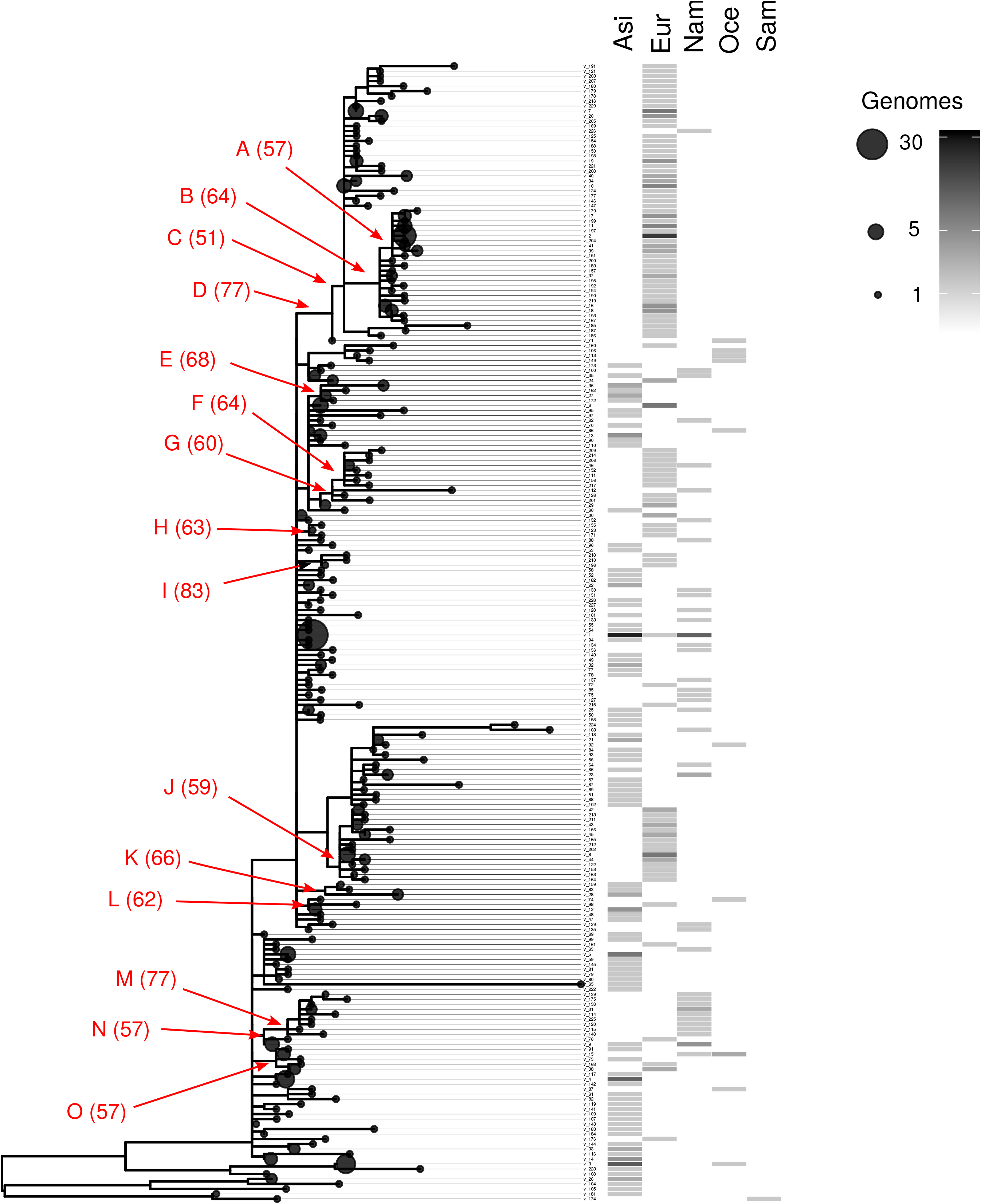
Phylogeny and distribution of 228 SARS-CoV-2 SVs corresponding to 353 genomes from the small dataset studied here. Dot sizes indicate the number of genomes represented by each SV, as indicated in the legend to the right of the Figure. Numbers in parenthesis indicate bootstrap supports of selected tree branches, provisionally named clades *A* to *O* for practical reasons. Tree terminals are aligned with the heatmap cells, which depicts the SVs’ distributions and abundances, as indicated by the scale to the right. *ASI Asia*; *EUR Europ*e; *NAM North America*; *OCE Oceania*; *SAM South Americ*a.

**Table 1.** Small dataset structure analysis.

|  | ntaxa<sup>1</sup> | obs<sup>2</sup> | rand.mean<sup>3</sup> | rand.sd<sup>4</sup> | obs.rank<sup>5</sup> | SES<sub>MPD</sub><sup>6</sup> | *p*-value<sup>7</sup> |
| --- | --- | --- | --- | --- | --- | --- | --- |
| Asia | 89 | 0.00048 | 0.00044 | 0.00008 | 8198 | 0.55351 | 0.81972 |
| Europe | 99 | 0.00031 | 0.00045 | 0.00006 | 1 | -2.28880 | 0.00010 |
| North America | 37 | 0.00027 | 0.00043 | 0.00009 | 19 | -1.80454 | 0.00190 |
| Oceania | 10 | 0.00043 | 0.00041 | 0.00014 | 6616 | 0.15601 | 0.66153 |

<sup>1</sup> Number of virotypes from each region (rows).
<sup>2</sup> Observed Mean Phylogenetic Distances (MPD).
<sup>3</sup> Average MPD in Monte Carlo (M-C) randomizations (dispersed model).
<sup>4</sup> Standard deviation of M-C MPDs.
<sup>5</sup> Observed MPDs ranks.
<sup>6</sup> Standardized effect of geographical structure.
<sup>7</sup> $H_0$ : evenly distributed virotypes.

**Figure 3.**
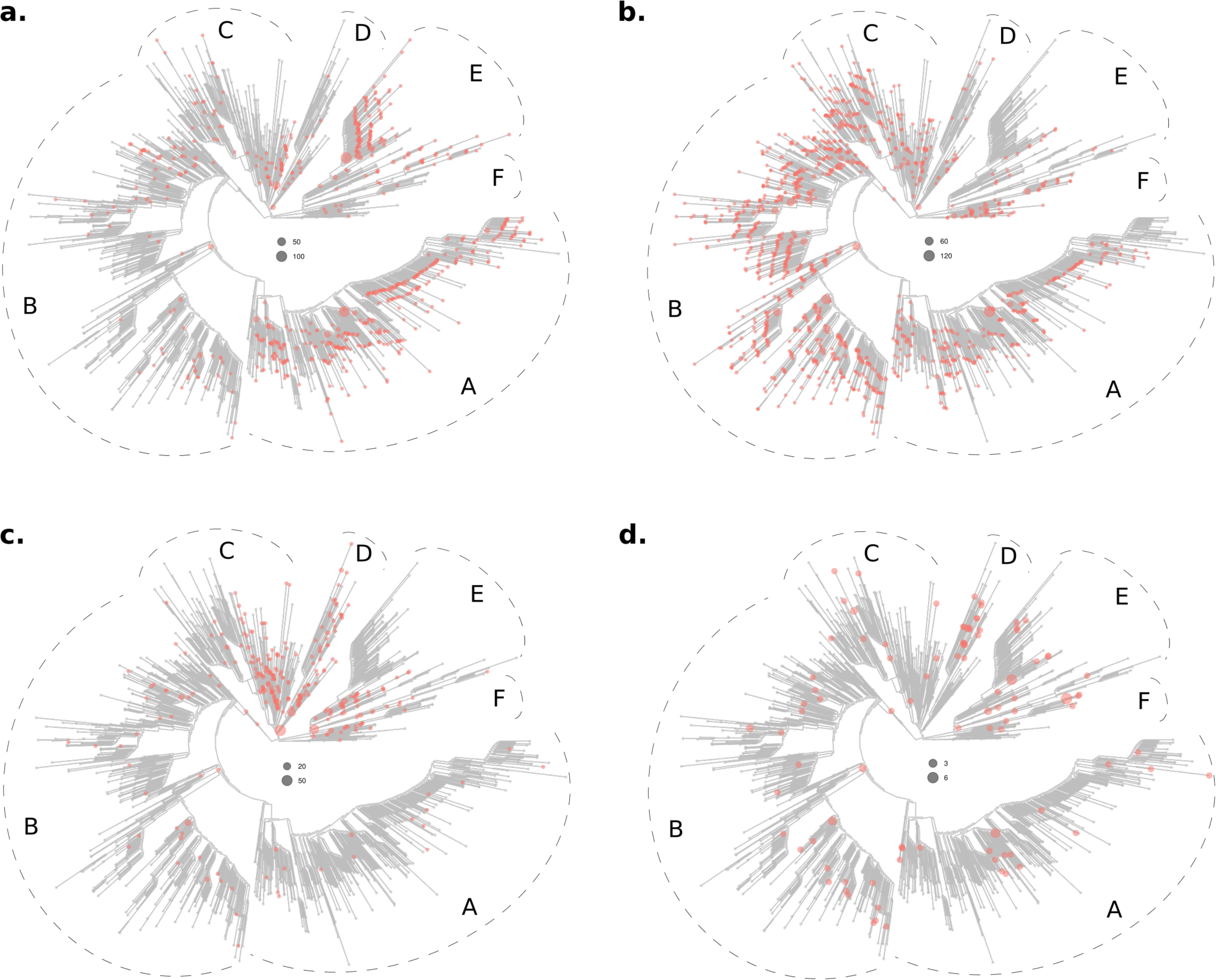
Phylogeny and distribution of 2,336 SARS-CoV-2 SVs representative of 4,333 genomes from around the world. The SVs present in in North America (**a**), Europe (**b**), Asia (**c**) and Oceania (**d**) are indicated by colored dots. Six sections of the tree are indicated (*A*, *B*, *C*, *D*, *E*, and *F*) to facilitate results interpretation (please see main text). Dots diameters are proportional to the number of genomes accrued in each SV, as indicated in each panel separately.

**Table 2.**
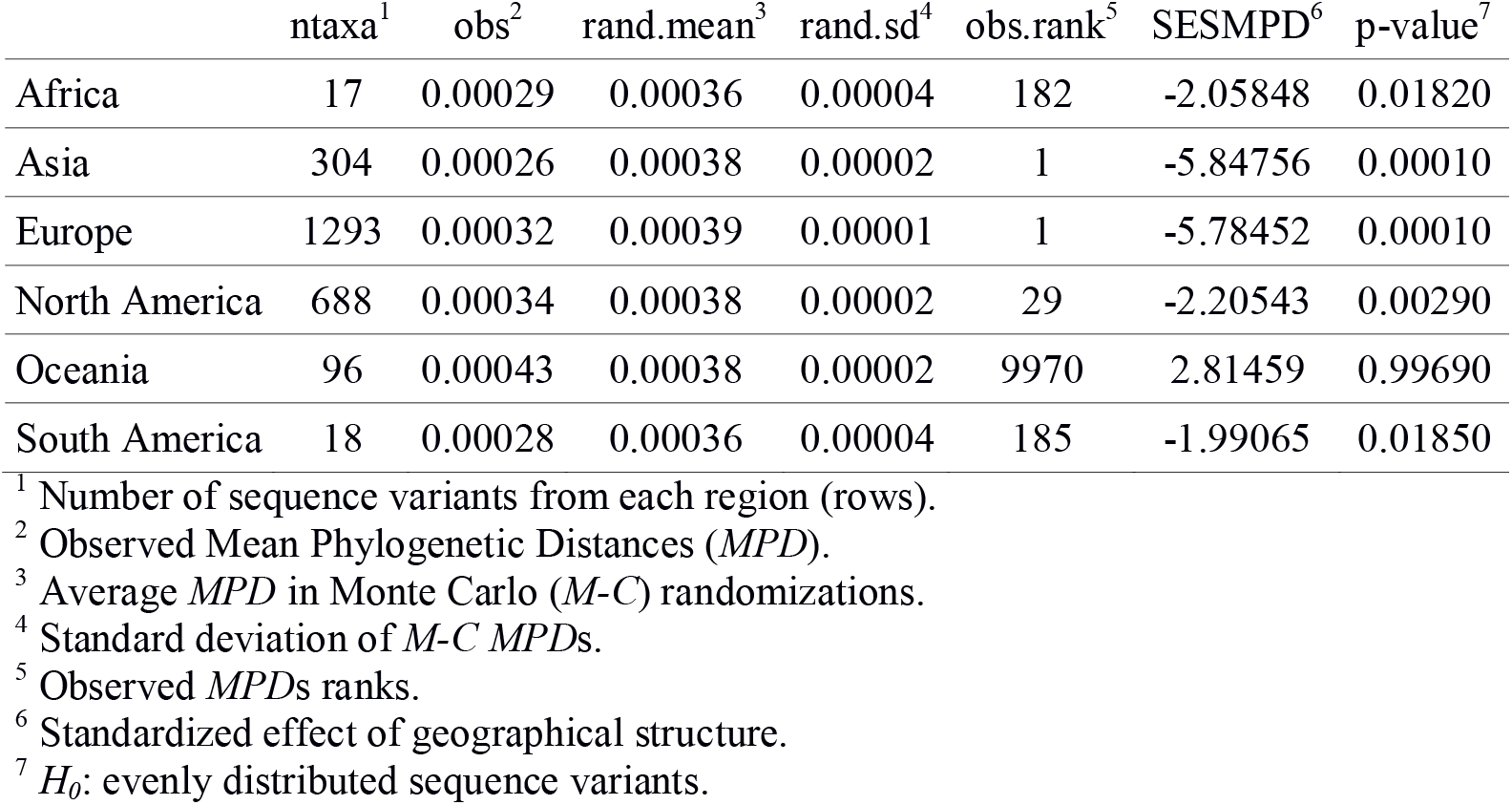
Large dataset structure analysis.

|  | ntaxa<sup>1</sup> | obs<sup>2</sup> | rand.mean<sup>3</sup> | rand.sd<sup>4</sup> | obs.rank<sup>5</sup> | SESMPD<sup>6</sup> | p-value<sup>7</sup> |
| --- | --- | --- | --- | --- | --- | --- | --- |
| Africa | 17 | 0.00029 | 0.00036 | 0.00004 | 182 | -2.05848 | 0.01820 |
| Asia | 304 | 0.00026 | 0.00038 | 0.00002 | 1 | -5.84756 | 0.00010 |
| Europe | 1293 | 0.00032 | 0.00039 | 0.00001 | 1 | -5.78452 | 0.00010 |
| North America | 688 | 0.00034 | 0.00038 | 0.00002 | 29 | -2.20543 | 0.00290 |
| Oceania | 96 | 0.00043 | 0.00038 | 0.00002 | 9970 | 2.81459 | 0.99690 |
| South America | 18 | 0.00028 | 0.00036 | 0.00004 | 185 | -1.99065 | 0.01850 |

<sup>1</sup> Number of sequence variants from each region (rows).
<sup>2</sup> Observed Mean Phylogenetic Distances (*MPD*).
<sup>3</sup> Average *MPD* in Monte Carlo (*M-C*) randomizations.
<sup>4</sup> Standard deviation of *M-C MPDs*.
<sup>5</sup> Observed *MPDs* ranks.
<sup>6</sup> Standardized effect of geographical structure.
<sup>7</sup> $H_0$ : evenly distributed sequence variants.

### Protein level phylogeographic analysis

To asses if the phylogenetic structuring process affected the virus proteins, we analyzed the distribution and ancestral trajectories of 11 amino acidic polymorphisms, all of which were endorsed by at least 100 sequences (Fig. 4). The geographic distributions of the corresponding amino acid variants were very heterogeneous. Furthermore, ancestral amino acidic transitions were highly fitted to the virus phylogeny and occurred at branches dividing the main tree sections described above, or separating large tree chunks inside these sections. These results are summarized in Figures 5–8. The distributions of the 57Q/H, 85T/I and 614 D/G polymorphisms of the orf3, nsp2 and S proteins, respectively, were very contrasting between the east and west (Fig 5). The amino acidic variants 57H, 85I, and 614G were frequent in Europe and North America, whereas 57Q, 85T, and 514D were relatively abundant in Asia. Unlike the majority of countries, China presented neither the 175M variant of the M protein nor amino acids K and R at positions 203 and 204, respectively, of the N protein (Fig 6). The nsp13 protein amino acids 504L and 541C were almost exclusively observed among the sequences from North America and Australia (Fig. 7 a and b). The 84L/S polymorphism (orf8 protein) displayed a similar geographic pattern. However, the 84S variant turned out to be abundant in China and South Korea as well as in North America and Australia (Fig. 7 c). The 37 and 251 positions of the nsp6 and orf3a proteins, respectively, experienced mutations inside the C tree section (Fig. 8). However, position 37 also experienced a change in tree section D. In agreement with this, the 215V variant was prevalent in Europe, China and Australia, whereas 37F was also abundant in Japan and relatively more abundant than 215V in Australia.

**Figure 4.**
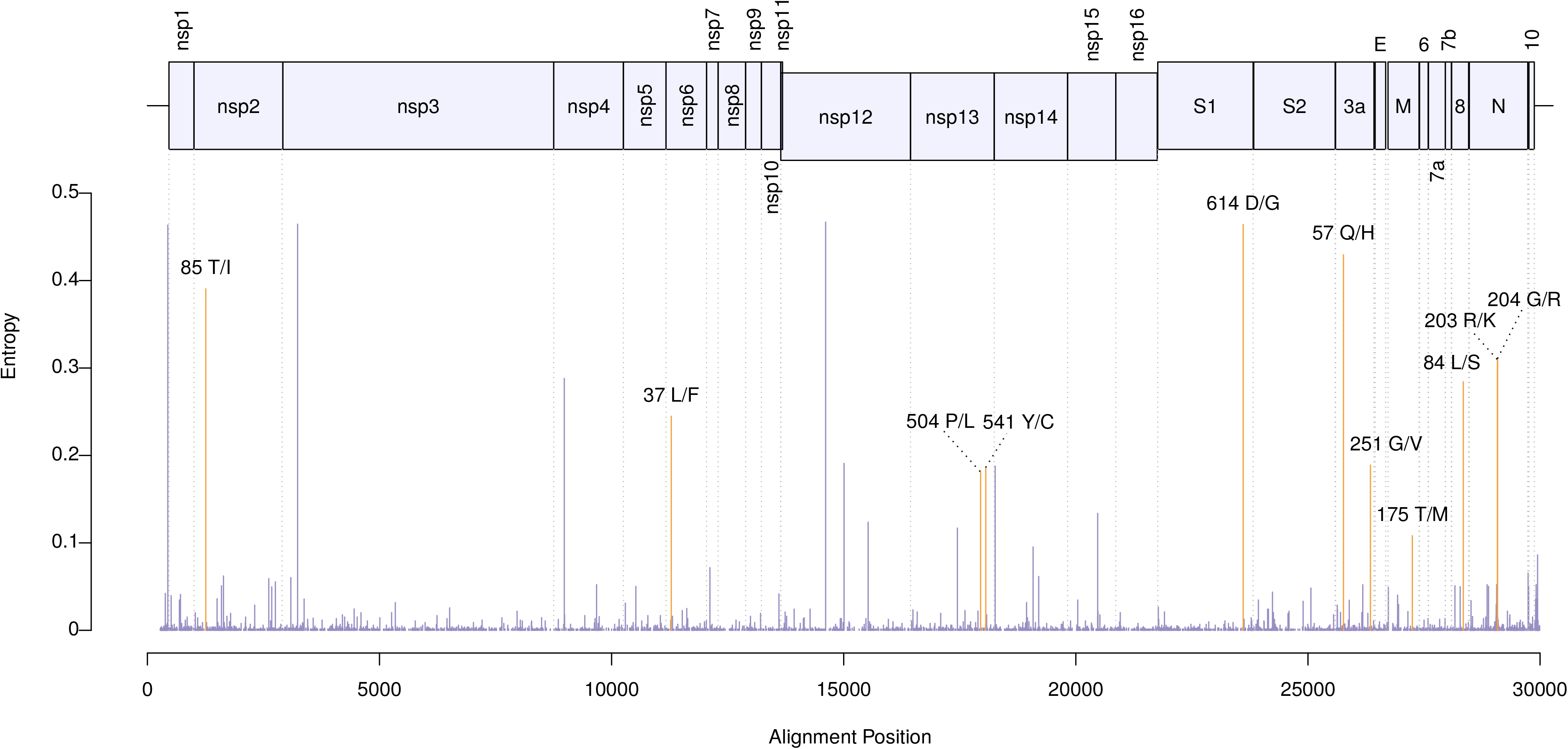
Location in the virus genome of 11 amino-acidic polymorphism used in phylogeographic analyses. The positional entropies (y-axis) were derived from the large dataset studied here. The orange-highlighted bars correspond to the positions that displayed non-synonymous mutations in at least 100 sequences.

**Figure 5.**
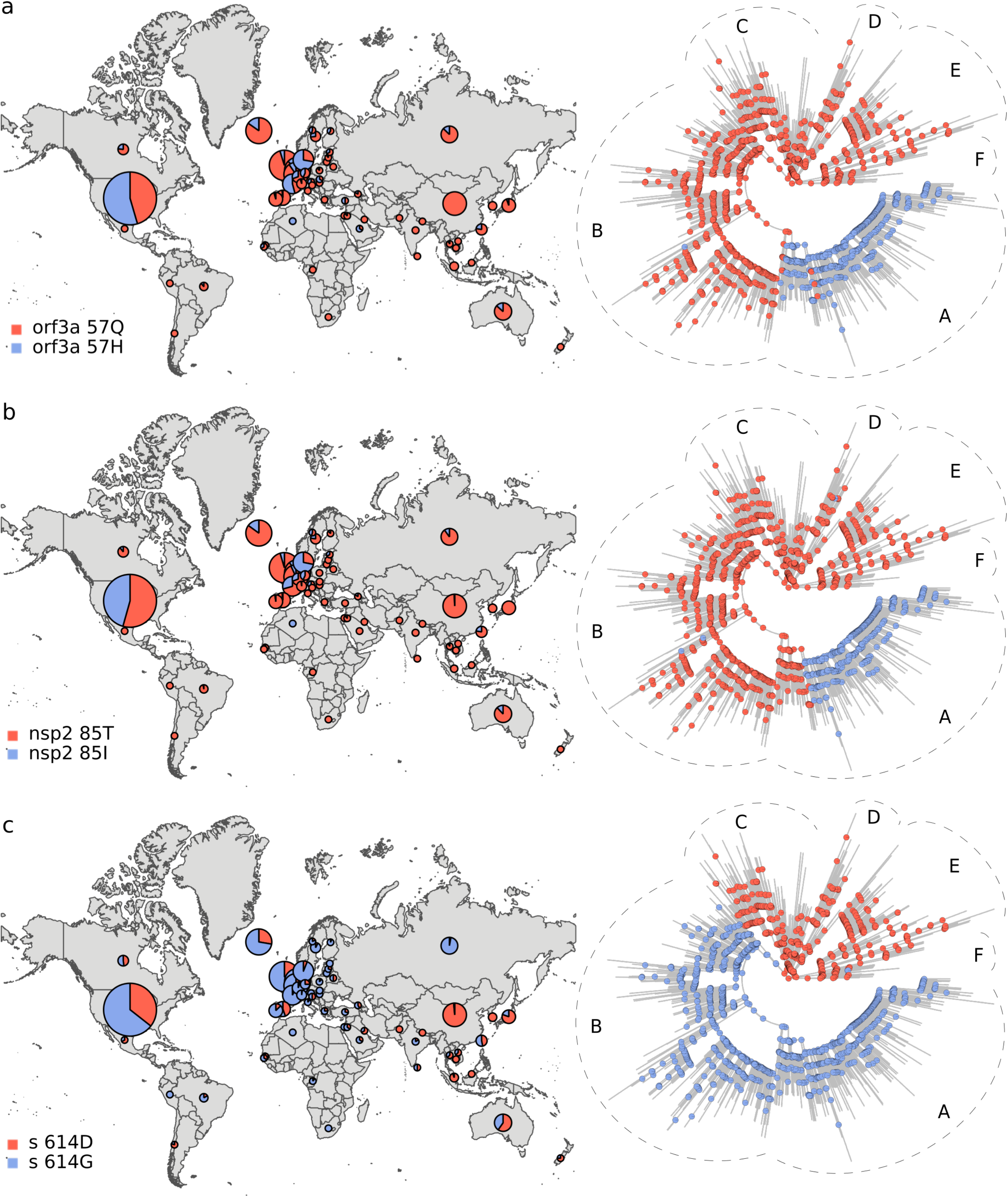
Geographic distribution and evolutionary trajectories of polymorphisms 57Q/H (a), 85T/I (b) and 614D/G (c).

**Figure 6.**
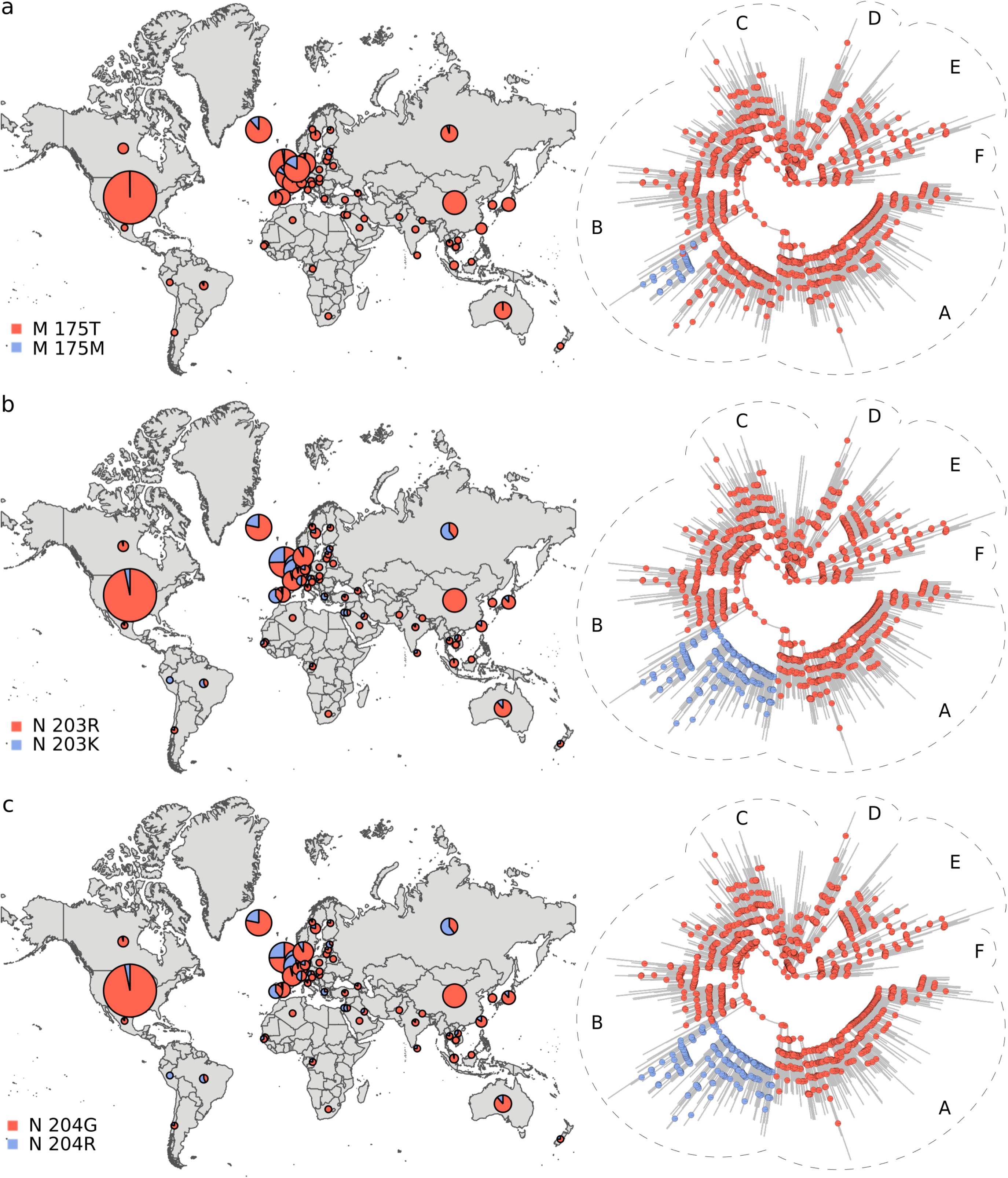
Distribution and evolutionary trajectories of polymorphisms 175T/M (a), 203R/K (b), and 204G/R.

**Figure 7.**
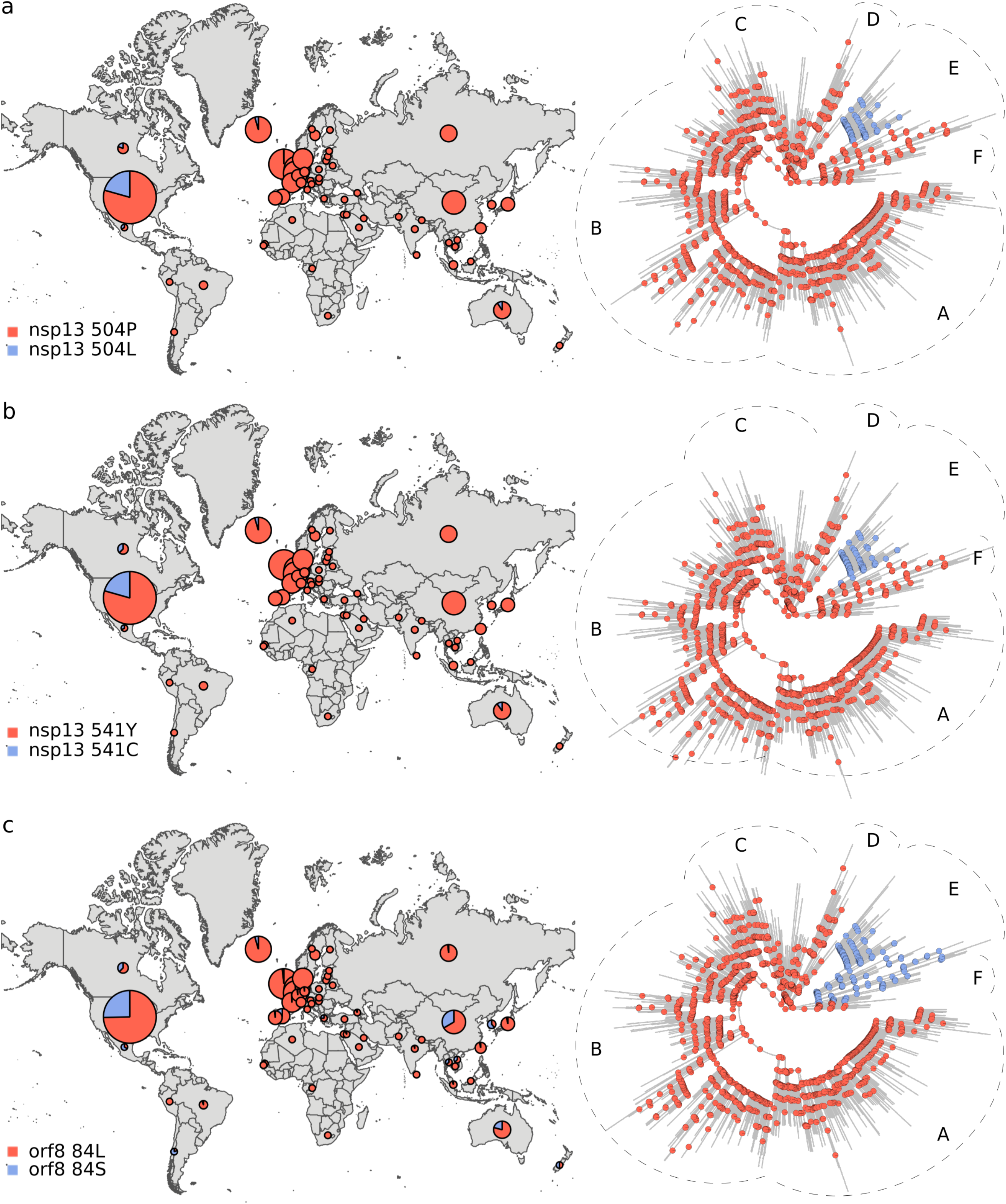
Distribution and evolutionary trajectories of polymorphisms 504P/L (a), 541Y/C (b), and 84L/S (c).

**Figure 8.**
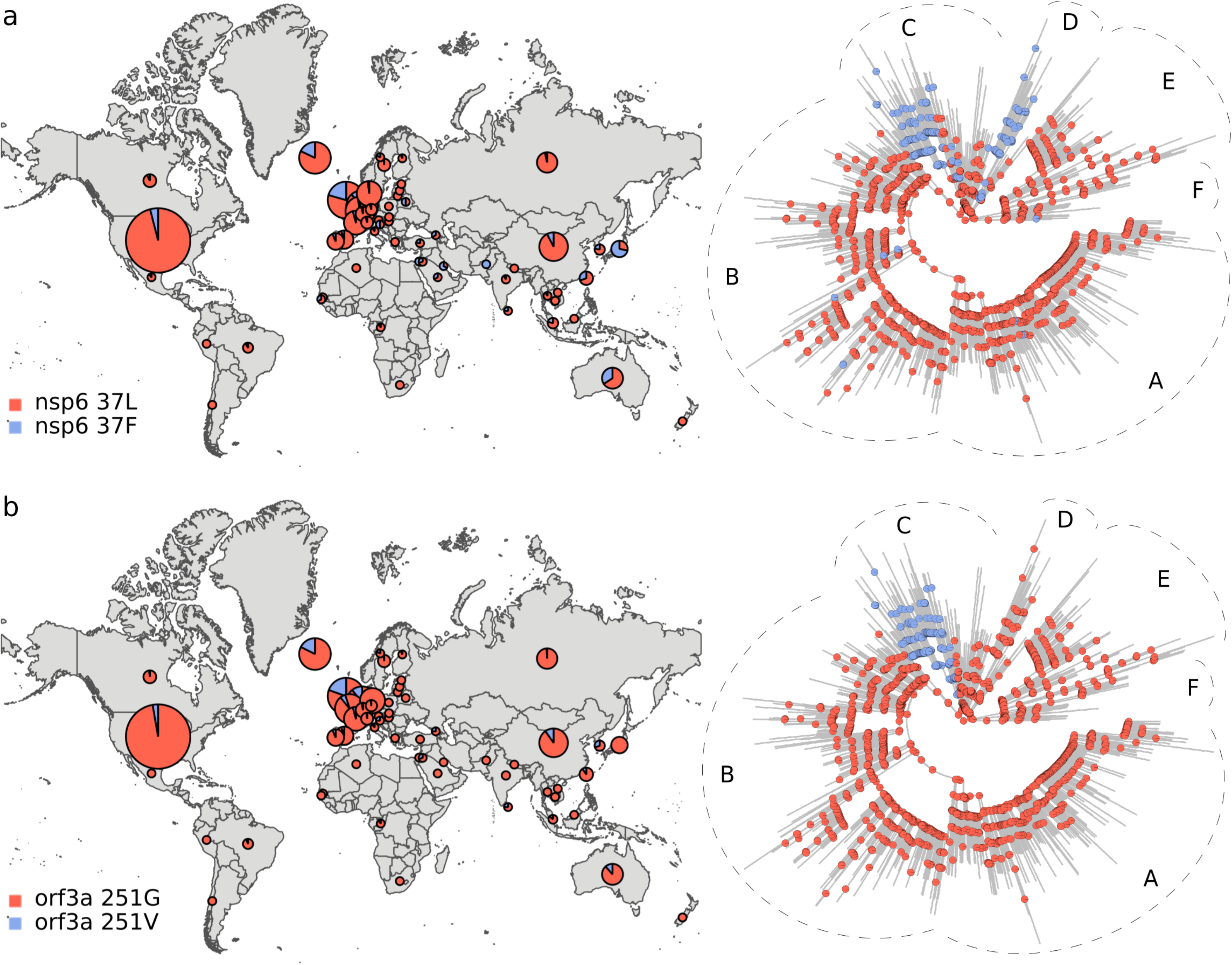
Distribution and evolutionary trajectories of polymorphisms 37L/F (a) and 251G/V (b).

## Discussion

### SARS-CoV-2 sequence variants

Using an exact algorithm, we identified 228 sequence variants among 353 SARS-CoV-2 genomes sampled from mid-December 2019 to March 14, 2020. In a similar way, pairwise comparisons made from a multiple alignment of 4,333 selected sequences sampled between December 2019 and April 25 2020, revealed 2,336 genetic variants. Less than half the here studied sequences were identical to at least one of the rest of sequences and, out of 9,385,278 genome pairs compared from the large dataset, only 50,839 showed no mutations. A raw extrapolation of our results suggests that, only among the ~60,000 SARS-CoV-2 genomes sequenced so far, there could be some 30,000 unique sequences. These data are overwhelming. We anticipate significant methodological challenges, especially if the number of SARS-CoV-2 sequences continues to increase at present rates. As explained in the Materials and Methods, the approach used here to analyze the small dataset is *O(N*^*2*^*L)* to *O(N*^*3*^*L)* complex. This means that, depending on the algorithm and software used, it would be very expensive to replicate the analyses for more than several hundred or some few thousand sequences. Likely impossible for more than 10,000 sequences (Blackshields et al., 2010). The alternative approach implemented here, that is to base sequence comparisons on a multiple sequence alignment (MSA), allows avoiding the costly pairwise alignment step. However, the approach also faces some challenges. MSAs usually present misaligned ends attributable to sequencing differences. This can be difficult to handle, especially for large datasets, since there is no standardized procedure to deal with the problem. The issue (which is not exclusive of SARS-CoV-2) is frequently addressed by trimming off the problematic alignment regions. However, this involves visual inspection and data dismissal, which will require automation and reproducible procedures. Standard procedures are neither available for dealing with incomplete sequences, which are sometimes padded with *N*s, and ambiguous base calls. Although in some cases it can be handy to simply discard such sequences, some incomplete sequences may contain useful information that is absent from other sequences.

### Spatial Phylogenetic Structure

Here, we presented statistical evidence that SARS-CoV-2 phylogeny is spatially structured. From a biological point of view, this imply that the virus gene pool varies by world region. Remarkably, the virus phylogenetic structure has a close correlate at the protein level. All ancestral nodes inside tree section A presented the 57H and 85I amino acidic variants (Fig. 5 a and b). This indicate that the divergence event leading to the split between tree section A and the rest of the phylogeny was accompanied by protein mutations in the orf3a and nsp2 proteins. The fact that the strains from tree section A were abundant in North America and Europe (Fig. 3 a and b), strongly suggests that viral dispersal to, or from, these regions was associated with the emergence of polymorphisms 57Q/H and 85T/I. In a similar way, the 614D/G, 504P/L, 541Y/C and 84L/S polymorphism may have resulted from the spread of the virus between the east and west (Figs. 5 c and 7 a-c, respectively). The 175T/M, 203R/K and 204G/R transitions occurred inside tree section B (Fig. 6), which was highly represented among the European sequences (Fig. 3 b). This, together with the presence of 203K and 204R in Russia, suggests that these polymorphisms may have emerged in Europe or Russia. In a similar way, the 37L/F and 251G/V polymorphisms likely emerged in association with dispersal to, or from, Europe (Fig. 8). Our ancestral states inferences indicated that the amino acidic position 37 of the nsp6 protein experienced several independent L/P transitions. Two such transitions occurred at the phylogeny branches separating tree sections C and D from the rest of the virus phylogeny (Fig. 8 a). These two tree sections were well represented in Europe and Japan, respectively, strongly suggesting that 37L/F emergence may has occurred twice as a consequence of independent dispersion events.

Biogeographic patterns have been previously observed in other coronaviruses (Chu et al., 2018) and very different viruses as phages and retroviruses (Díaz-Muñoz et al., 2013; Rodriguez et al., 2009). That SARS-CoV-2 has developed a biogeography despite its high propagation rate may seem contradictory. However, spatial diversification depends not only on dispersion constraints but also on evolutionary rates. In particular, spatially structured phylogenies can be the consequence of speciation rates being very high relative to dispersion rates (Webb et al., 2002). As far as we know, travels between remote places constitute the only dispersal mechanism of the virus. This implies that founder viruses usually carry very small fractions of the total genetic variation of the source populations. Therefore, each time the virus spreads, substantial losses of diversity can occur, possibly combined with rare mutations settlement, due to founder effects. In addition, it stands to reason that, after dispersal, mutations accumulate quickly as newly established populations enlarge, due to viral polymerases high error rates (Gago et al., 2009; Moya et al., 2004). Some of these mutations can lead to novel phenotypes, as shown in Figures 3–8. Based on these considerations, it is reasonable to hypothesize that long-distance dispersal constitutes an opportunity for the virus to fix otherwise rare, and/or develop new, mutations.

The 2002-2003 SARS-CoV epidemic was subdivided into three genetically different phases, indicating that coronaviruses can mutate recurrently along relatively short periods of time (The Chinese SARS Molecular Epidemiology Consortium, 2004). It has been shown that slight mutations can produce significant phenotypic effects in MERS-CoV and other coronaviruses (Chu et al., 2018; Rasschaert et al., 1990; Zhang et al., 2007). Furthermore, recent experimental studies suggest that the 614G spike protein variant increases SARS-CoV-2 infectivity (Korber et al., 2020; Zhang et al., 2020). Here we show that at least 10 further protein polymorphisms may have emerged along the recent SARS-CoV-2 evolutionary history. This must be taken as a call for attention. The virus evolution should continue to be monitored and relationships should be sought between viral diversity and pathogenesis.

## Acknowledgments

We acknowledge the authors, originating and submitting laboratories of the sequences from GISAID’s EpiCoV™ Database used in this study. This work was partially supported by PIP 11220130100255CO (CONICET), PICT 2016-2795 (ANPCyT) and Asociación Civil Argentina Genetics.

## Notes

### Competing Interest Statement

The authors have declared no competing interest.

### Summary of Updates

This version includes new analyses based on protein sequences.

