## Supplementary figure legends for "Assessing SARS-CoV-2 spatial phylogenetic structure: Evidence from RNA and protein sequences"

**Figure S1.** Toy structured (***a***) and random (***b***) distributions. The colored squares represent four hypothetical areas (*A*, *B*, *C* and *D*). Colored branches correspond to evolutionary distances between region *B* lineages. Notice the smaller distances in the spatially structured phylogeny.

**Figure S2.** Diversity profile of selected SARS-CoV-2 genomes studied here (Materials and Methods). The entropies of the UTR alignment positions that have an abnormally high entropy are displayed by black bars.

**Figure S3.** Phylogeny and distribution of 2,336 *SARS-CoV-2* SVs corresponding to 4,333 genomes from Africa (*AFR*), Asia (*ASI*), Central America (*CAM*), Europe (*EUR*), North America (*NAM*), Oceania (*OCE*) and South America (*SAM*). Internal nodes are labeled with dots colored according to the corresponding bootstrap supports, as indicated by the scale to the right. Tree terminals are aligned with the cells of the heat map, which depicts the SVs’ distributions and abundances, as indicated by the scale to the right.
