## Supplementary figures and images for "Assessing SARS-CoV-2 spatial phylogenetic structure: Evidence from RNA and protein sequences"

### Figure S1

a

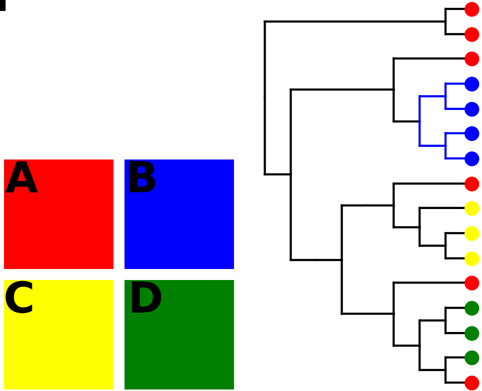


b

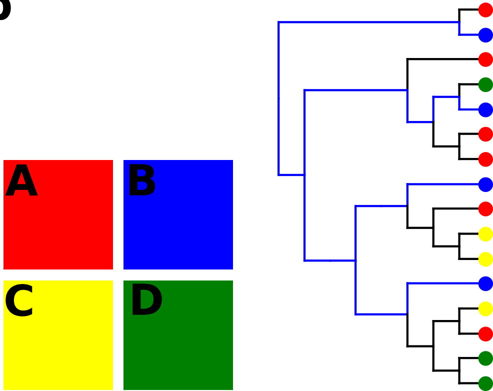

### Figure S2

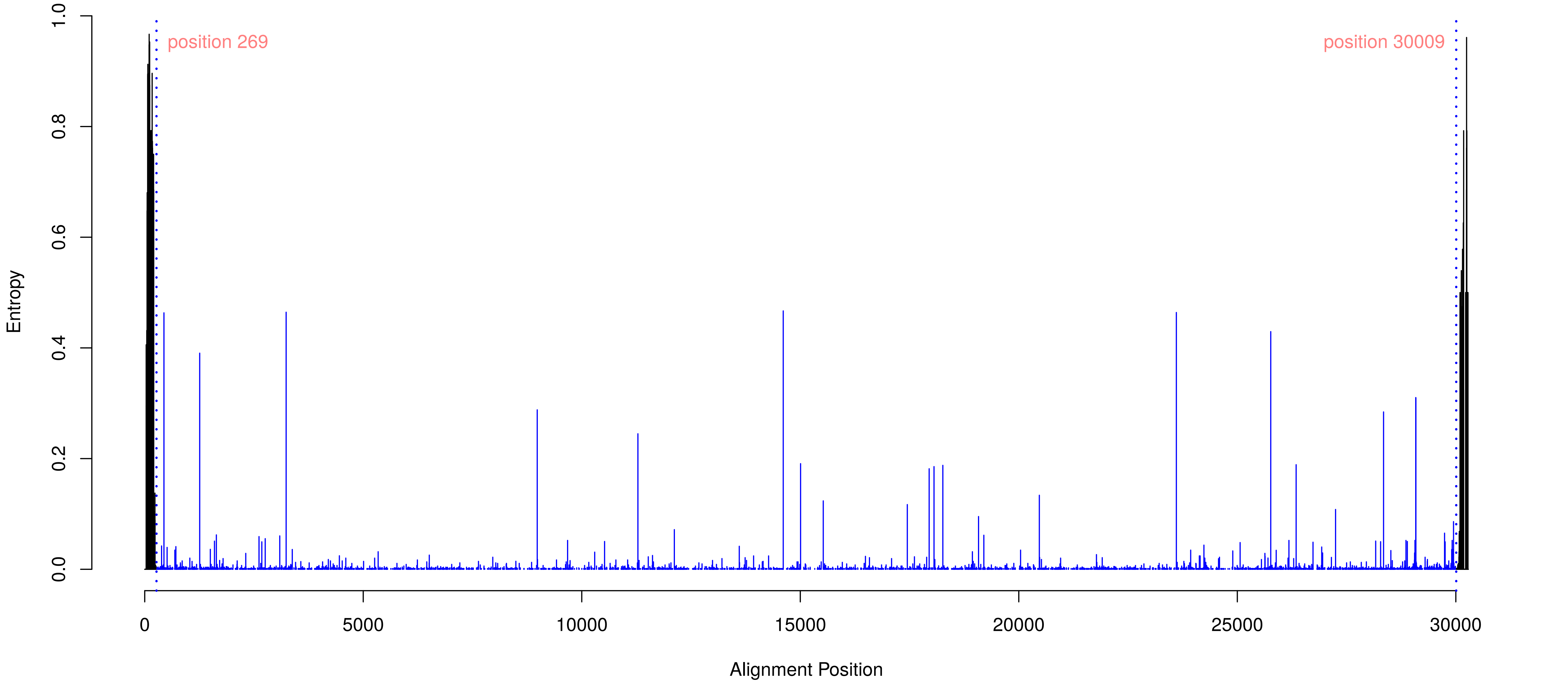

### Figure S3

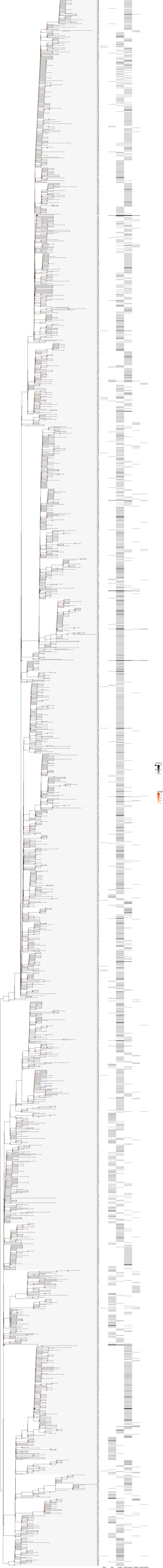
